# Inter- and intra-individual differences in β-band sensorimotor rhythm modulation and its relation to corticomuscular coherence in force-tracking tasks

**DOI:** 10.1101/2025.11.10.687531

**Authors:** Jing Zhou, Junichi Ushiyama

**Author notes:** **Author Contributions**: Conceptualization, JZ and JU; Methodology, JZ and JU; Data acquisition, JZ; Data Analyses, JZ; Visualization, JZ; Interpretation, JZ and JU.; Writing – Original Draft, JZ; Writing – Review & Editing, JZ and JU; Supervision, JU; Funding Acquisition, JZ and JU. **Conflict of interest**: The authors declare no competing interests. **Code availability**: The computer code that supports the findings of this study is publicly available on GitHub at https://github.com/JingZhou-95/Time-resolved-modulation-of-beta-band-sensorimotor-rhythm.git. **Data availability**: Source data for this study are not publicly available due to ethical restrictions. The source data are available to verified researchers upon reasonable request by contacting the corresponding author.

## Abstract

The β-band sensorimotor rhythm (SMR), recorded using electroencephalography, generally begins to decrease in amplitude with motor preparation reflecting cortical desynchronization, and subsequently becomes coupled with electromyogram signals during voluntary contraction, thus forming corticomuscular coherence (CMC). However, it remains unclear how the time-resolved modulation of β-band SMR varies among individuals, potentially relate to individual differences in CMC. Additionally, it is also unclear how the nervous system modulates β-band SMR to meet varying task demands within individuals. Here, we explored how time-resolved modulation patterns of β-band SMR relate to CMC from two perspectives: inter-individual differences (Experiment 1), and intra-individual variability depending on task demands (Experiment 2). In Experiment 1, participants repeated a steady-force maintenance task. The degree of β-band SMR modulation (i.e., rebound from desynchronization to synchronization) during contractions varied greatly among individuals and was positively correlated with CMC magnitude. This suggests that even under steady-force maintenance, the motor control strategy used while regulating β-band SMR varies greatly among individuals. In Experiment 2, participants who showed significant CMC in Experiment 1 performed four tasks with varying target trajectories. Even within individuals, the degree of β-band SMR modulation was reduced in parallel with the CMC magnitude as force adjustment was required. This finding suggests that, when confronted with more challenging task demands, our nervous system suppresses the SMR modulation and decouples from muscles. Overall, the way in which our nervous system regulates β-band SMR is assumed to represent a strategy for flexible adaptation to diverse motor environments.

**NEW & NOTEWORTHY:** This study is the first to track how corticomuscular coherence (CMC) and β-band sensorimotor rhythm (SMR) co-vary during voluntary isometric contraction, from both inter- and intra-individual perspectives. Across individuals, CMC magnitude was tightly related to the degree of β-band SMR modulation (rebound from desynchronization to synchronization). Within individuals, β-band SMR modulation and CMC were changed in parallel with task demands. Regulation of β-band SMR may serve as a basis for flexible motor control.

## Introduction

The sensorimotor rhythm (SMR) refers to rhythmic, synchronized neural activity that is recorded over the sensorimotor cortex (Lopes da Silva, 1991). The β-band (15–35 Hz) SMR modulations has been considered to be implicated in sensorimotor processing during motor-related events, including motor preparation, motor execution, and post-movement processes, highlighting their critical role in motor control (Pfurtscheller & Lopes da Silva, 1999; Brovelli *et al*., 2004).

A power reduction of the SMR within the β-bands around motor initiation, known as movement-related β-band desynchronization (MRBD) (Espenhahn *et al*., 2017a; Zhang *et al*., 2025) is caused by desynchronization of the neurons within the sensorimotor area as the result of voluntary movement or passive movement (Pfurtscheller, 1977; Cassim *et al*., 2001; Adham *et al*., 2024). After movement offset, β-band SMR power shows a transient increase, known as the post-movement β-power rebound (PMBR) (Parkes *et al*., 2006a; Fry *et al*., 2016). Moreover, β-band SMR power also tends to increase once the contraction or posture becomes stable (Cassim *et al*., 2000; Alegre *et al*., 2006; Spinks *et al*., 2008; Van Elk *et al*., 2010). To explicitly distinguish it from the PMBR, we operationally define this power increase during steady contraction as steady-related β-band synchronization (SRBS). While MRBD is interpreted as reflecting an activated sensorimotor cortical state (Neuper & Pfurtscheller, 2001; Yuan *et al*., 2010), SRBS during steady contraction is suggested to signal the need to maintain the current motor set (Engel & Fries, 2010; Spitzer & Haegens, 2017).

Corticomuscular coherence (CMC) represents the synchronization between the cortex and muscle during voluntary muscle contraction. Indeed, the electroencephalogram (EEG) signals over the sensorimotor cortex are reported to be significantly coherent with the electromyogram (EMG) signals recorded from the contracted muscle within the β-band during voluntary contractions (Conway *et al*., 1995; Mima *et al*., 2002; Kristeva *et al*., 2007; Chakarov *et al*., 2009; Peng *et al*., 2024). Coherence function itself can be mathematically interpreted as the frequency-domain analogue of the squared correlation coefficient between two signals and is theoretically independent of the absolute power of either signal (Fancourt & Parra, 2001; Julius S. Bendat, 2010). However, CMC magnitude has been shown to correlate positively not only with SRBS (Kristeva *et al*., 2007; Ushiyama *et al*., 2011), but also with MRBD magnitude (Bayraktaroglu *et al*., 2013; Sun *et al*., 2025). These associations may appear paradoxical because CMC is linked to β-power changes in opposite directions (i.e., both increases and decreases). If both associations valid, we assume that the significant factor relates to CMC is not a unidirectional β-power change per se, but rather the overall modulation profile of the β-band SMR during motor control (i.e., the modulation span from MRBD to SRBS). We believe this approach to be highly valuable because no studies have investigated how CMC and β-band SMR change in relation to one another as the task progresses. Here, we conducted two experiments to examine how CMC relates to β-band SMR modulation over time. Experiment 1 involved a steady-force maintenance task to identify inter-individual differences. Participants who exhibited significant CMC in Experiment 1 proceeded to Experiment 2, which included a steady-force maintenance task and three types of dynamic-force adjustment tasks with varying task difficulties to explore intra-individual differences. This within-individual approach will allow us to observe how the time-resolved covariation between β-band SMR modulation and CMC adapts across conditions, thus advancing our understanding of the physiological mechanisms underlying how coordination between cortical and muscular activity changes depending on task difficulty. Using this two-tiered experimental design, the present study uniquely integrates inter- and intra-individual results to provide a robust understanding of the time-resolved interaction patterns between β-band SMR modulation and CMC, offering detailed insights into the neural activity that supports adaptive motor control in humans.

## Materials and Methods

### Participants

We recruited 30 healthy young Asian adults (19 males and 11 females, age range 20– 31 years) to participate in Experiment 1. Of these, nine participants exhibited significant CMC and proceeded to participate in Experiment 2. All participants were right-handed and none reported a history of neuromuscular or musculoskeletal disorders. Each participant provided informed consent prior to participation. The experiments were performed in accordance with the Declaration of Helsinki and were approved by the SFC Research Ethics committee in Shonan Fujisawa Campus, Keio University (Approval Number 167).

### Recordings

An ankle dynamometer (TCF100N; Takei Scientific Instruments Co., Ltd., Niigata, Japan) was fitted to each participant’s right foot via two straps to record their force in real time and reflect it as visual feedback. The force signal was low-pass filtered using a second-order Butterworth filter at a cutoff frequency of 50 Hz and was amplified using an amplifier (DPM-711B; Kyowa Electronic Instruments Co., Ltd., Tokyo, Japan). Scalp EEG signals were recorded from 29 cap-mounted (g.GAMMAcap 1027; Guger Technologies, Graz, Austria) Ag/AgCl electrodes (diameter of 18 mm, g.LADYbirdPASSIVE 1035, Guger Technologies), which were arranged according to the 10–20 international system (NeuroScan, Inc., Herndon, VA, USA) (Fig. 1A). Reference and ground electrodes were placed on the right and left earlobes, respectively. The EEG signals were amplified and band-pass filtered between 0.5–200 Hz using an analog biosignal amplifier (g.BSamp 0201A; Guger Technologies). Surface EMG signal was recorded over the muscle belly of the right tibialis anterior (TA) muscle using bipolar Ag/AgCl electrodes with a diameter of 10 mm and an interelectrode distance of 30 mm. The EMG signals were amplified and band-pass filtered between 5–500 Hz (g.BSamp 0201A; Guger Technologies).

The analog force, EEG, and EMG signals were converted into digital signals by an analog-to-digital converter (NI cDAQ-9178 with NI 9215 modules, National Instruments, Austin, TX, USA) at a sample rate of 1000 Hz; it was controlled by a datalogger program that was originally designed using MATLAB software (MathWorks, Inc., Natick, MA, USA).

### Experimental Design

Participants were seated comfortably in a chair with the knee flexed at 90° and the ankle extended at 10°. Straps were used to secure the knee and foot to maintain a stable posture. A monitor was located 2 m in front of participants at eye level. It displayed the motor tasks and visual feedback (Fig. 1B). Isometric contractions of the TA muscle controlled a red square marker on the screen, which moved up or down according to the force that was exerted. Prior to the tasks, we measured each participant’s maximal voluntary contraction (MVC) force for ankle dorsiflexion after sufficient practice to become familiar with activating the TA muscle as needed.

In the task period, each trial consisted of five phases: “Rest,” “Relax,” “Ready,” “Contraction,” and a final “Relax.” During the 7-s “Rest” phase, participants were allowed to move freely to adjust their posture. Participants were then required to remain stationary during the following 5-s “Relax” phase. In the subsequent 3-s “Ready” phase, three auditory cues (frequency, 500 Hz; duration, 80 ms) were presented at 1-s intervals to prepare participants for the upcoming motor task. In the 7-s “Contraction” phase, participants were instructed to initially perform ballistic contraction by quickly dorsiflexing their right ankle (force overshoot relative to the target line frequently occurred), and then promptly adjust the force to align the marker with the target line as precisely as possible. After completing the contraction, participants immediately ceased contracting and entered a final 3-s “Relax” phase, during which they remained stationary and avoided further muscle activity. Visual phase labels in combination with a brief auditory cue (frequency, 1000 Hz; duration, 300 ms) were presented at the transition of each phase to guide participants. Specifically, if exaggerated postural adjustment or verbalization occurred during relax, ready and contraction phase, the trial was considered invalid and an additional trial was conducted.

In Experiment 1, the target lines were simple straight lines set at heights corresponding to 10% (10%-task) and 15% (15%-task) of the MVC. Each force level was performed in two blocks of 16 trials each (32 trials per task). All trials within a block involved only one task type to avoid mixing task conditions, and the order of the blocks was randomized. Participants were given sufficient rest (5–10 minutes) between blocks to prevent fatigue.

Experiment 2 introduced more complex target line patterns (Fig. 1C), as follows: 1) a straight line at 10% MVC (St-task, which was the same as the 10%-task in Experiment 1); 2) a single sine wave with a period of 7 s (1sin-task); 3) three consecutive sine waves with a period of 2.3 s each (3sin-task); and 4) a composite curve made up of four sine waves with periods of 2.3 s, 1.2 s, 2.3 s, and 1.2 s (4sin-task). All sinusoidal waveforms had an amplitude of 2% MVC and an average height of 10% MVC, and task difficulty was manipulated by increasing the rate of force adjustment and introducing variability in that rate over time. Figure 1D illustrates the time course of the target force derivative, i.e., the required force-change rate, for the three dynamic force-adjustment tasks. The 1sin-task required tracking a relatively slow target modulation, whereas the 3sin-task increased difficulty by requiring more frequent and faster adjustments. The 4sin-task imposed a variable rate demand in which the target force derivative alternated between slower and faster segments over time, so participants had to switch between different rate requirements. Each task type was conducted in two blocks of 16 trials each (32 trials per task); all trials within a block involved the same task, and the order of the blocks was randomized. Sufficient rest (5–10 minutes) was provided between blocks based on each participant’s needs.

**Figure 1:**
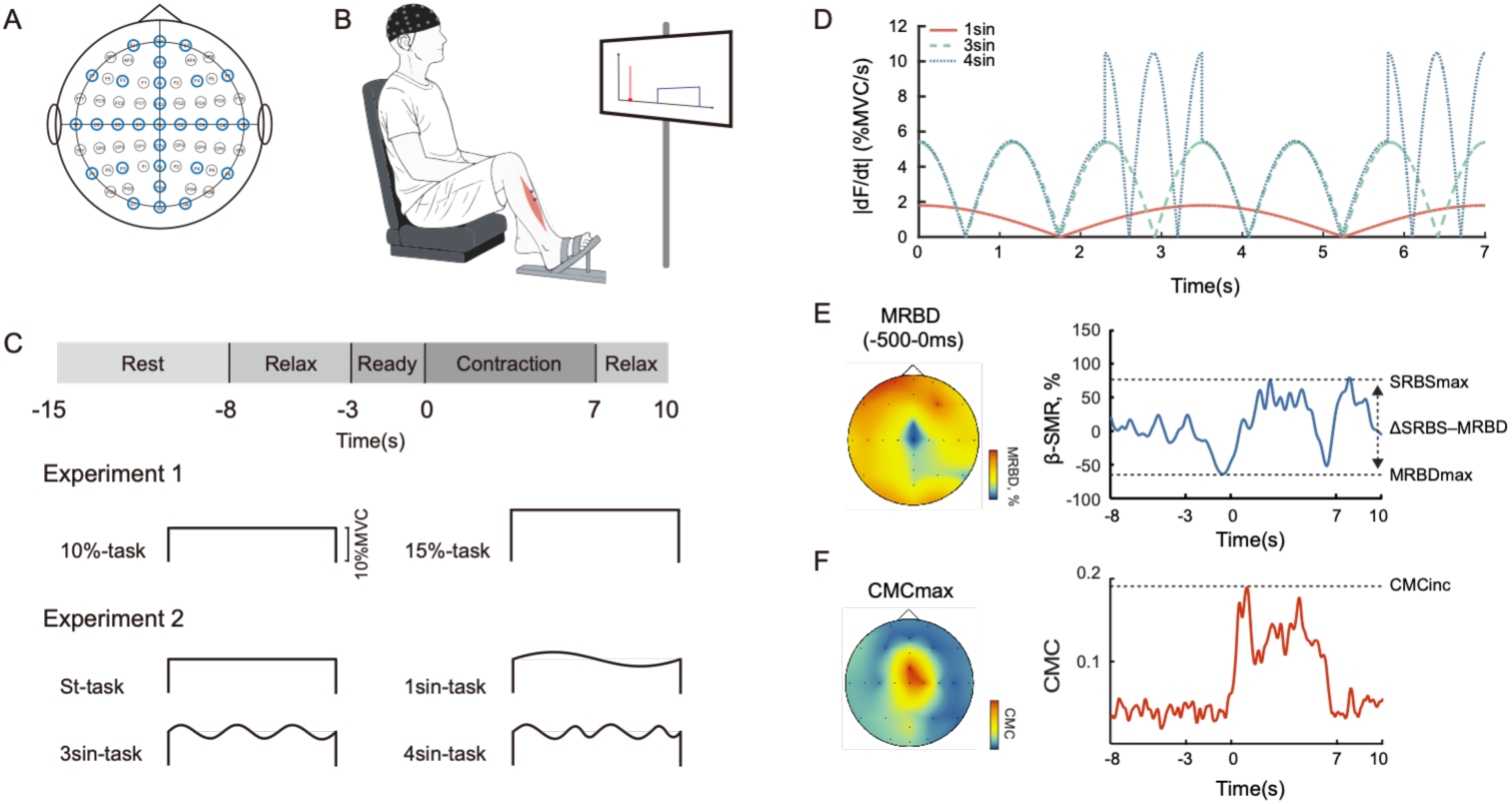
Research methods for Experiments 1 and 2. (A) EEG electrode layout. EEG was recorded from 29 scalp channels (blue). (B) Schematic overview of the experimental setup. Participants were seated comfortably while EEG, EMG from the tibialis anterior muscle, and right ankle force were recorded. Real-time visual feedback of the exerted ankle dorsiflexion force was provided for the visuo-motor tasks. (C) Motor tasks. Each trial had five phases: 7-s “Rest,” 5-s “’Relax,” 3-s “Ready,” 7-s “Contraction,” and 3-s “Relax.” In Experiment 1, straight target lines were set at 10% and 15% of the maximal voluntary contraction (MVC). In Experiment 2, the following tasks were included: 1) “St-task” showing a straight line at 10% MVC, 2) “1sin-task” showing a sine wave with a 7-s period, 3) “3sin-task” showing three connected sine waves with a 2.3-s period, and 4) “4sin-task” showing a composite curve of four sine waves with periods of 2.3 s, 1.2 s, 2.3 s, and 1.2 s. All sine waves had a 2% MVC amplitude and averaged 10% MVC. Participants were instructed to control their muscle contraction to trace the target line as accurately as possible during the contraction phase. (D) Time course of the required target force derivative, i.e., force-change rate in the dynamic force-adjustment tasks of Experiment 2. Solid red, dashed green, and dotted blue lines represent the 1sin-, 3sin-, and 4sin-tasks, respectively. (E) Left: Topographic distributions of MRBD (−500 to 0 ms) from a typical participant during the 10%-task. Blue indicates greater MRBD. Right: Representative time course of β-band SMR responses during the motor task. MRBDmax was defined as the minimum value, and SRBSmax as the maximal value of EEG β-power during contraction relative to the baseline. The ΔSRBS–MRBD was calculated as the difference between SRBSmax and MRBDmax. (F) Left: Topographic distributions of maximal β-band CMCmax from a typical participant during the 10%-task. Red indicates stronger CMC. Right: Representative time course of β-band CMC during the motor task. CMCinc was defined as the maximal value of the β-band CMC within the contraction period (0–7 s), as indicated by the dashed line. Both CMC and MRBD exhibited focal distributions centered over the medial primary motor cortex, corresponding to lower limb representation.

### Data Analyses

#### Data Preprocessing

The EEG and EMG data were preprocessed using MATLAB (R2023a) and EEGLAB (v.2022.0) (Delorme & Makeig, 2004). Trials in which participants exhibited behaviors that may potentially affect the signals, such as speaking, moving, or failing to complete the trial, were excluded. The number of valid trials per tasks per condition varied across participants (Experiment 1: 10%-task, 26.7 ± 3.4; 15%-task, 28.5 ± 4.3; Experiment 2: St-task, 29.4 ± 2.7; 1sin-task, 30.0 ± 2.8; 3sin-task, 30.1 ± 2.7; 4sin-task, 29.1 ± 3.1). To maintain equal trial counts across tasks while maximizing participant inclusion, 24 trials were randomly selected per task for each participant. Participants with fewer than 24 trials in any task category after artifact rejection were excluded from group-level comparisons. Accordingly, one participant was excluded from the 10%-task and another from the 15%-task in Experiment 1 due to insufficient valid trials. Additionally, one participant in the 10%-task and two in the 15%-task were excluded due to data errors. No participants were excluded in Experiment 2.

For all tasks, data from the “Rest” phase were excluded, and the “Relax,” “Ready,” and “Contraction” phases (18 s in total) remained for analysis. All EEG and EMG signals were notch-filtered at 50 Hz (bandwidth 49–51 Hz) to remove electrical power noise. EMG data were rectified (rEMG) to emphasize the grouped discharge of motor units and extract the oscillatory envelope, which is considered advantageous for CMC analysis especially at low contraction levels (Farina *et al*., 2013; Ward *et al*., 2013; Dakin *et al*., 2014). EEG data were band-pass filtered between 1–100 Hz. EEG artifacts and non-brain signals were removed using Independent Component Analysis with the runica algorithm in EEGLAB (Bell & Sejnowski, 1995). ICA components corresponding to eye blinks and eye movements were visually identified based on their characteristic time courses and then rejected (Delorme *et al*., 2007).

Because the main procedures for MRBD/SRBS and CMC calculations are described in the “MRBD and SRBS analysis” and “CMC analysis” sections, here we briefly outline the topographical mapping steps to investigate the global spatial distribution of MRBD and CMC. The EEG data were first re-referenced to the average by subtracting the mean amplitude across all channels from each channel (Osselton, 1965). For MRBD, the time-resolved MRBD was obtained for each channel, and the topographical map was constructed using the mean MRBD within the −500 ms to 0 ms window relative to the onset of the “Contraction” phase (Fig. 1E, left). For CMC, we computed the CMC spectrum for each EEG channel and extracted the maximal CMC value of the contraction phase (2–6 s) to generate the CMC topographical map (Fig. 1F, left). On the basis of these spatial distributions, we confirmed that both MRBD and CMC were most pronounced around the Cz channel (over the sensorimotor area of the lower limb). In the next step of the subsequent main analysis, we specifically focused on the Cz channel, and applied a small Laplacian filter to the Cz channel by subtracting the averaged potentials from the surrounding channels (C1, C2, FCz, and CPz) to emphasize the EEG signal for CMC and MRBD analyses (Hjorth, 1975).

#### MRBD and SRBS analysis

EEG data from each trial were segmented into 1-s epochs with 50-ms sliding steps. For each time point, the 1-s epochs from all 24 trials within the same task condition were concatenated for power spectral density (PSD) estimation using Welch’s method (Hanning window; overlap, 0). This procedure produced 360 PSD estimates across an 18-s task period (excluding the 7-s “Rest” phase), based on a data matrix of (1000 data points/window × 360 windows × 24 trials), which were assembled into an EEG time–frequency analysis. MRBD and SRBS were then computed from concatenation-based PSD estimates (Meirovitch *et al*., 2015) using the following equation:

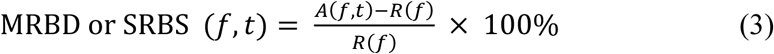

where *A* is the EEG PSD at time *t* and frequency *f*, and *R* is the mean PSD of the baseline period (the last 1 s in the 5-s “Relax” phase). Negative values reflect MRBD whereas positive values reflect SRBS. More negative MRBD values indicate greater EEG PSD suppression during the task. The frequency band of interest was in the broadband β-range (15–35 Hz), which covered the main frequency range of CMC, to investigate how β-band SMR modulation relates to CMC. The magnitude of MRBD (MRBDmax) was quantified by identifying the maximum percentage decrease in β-range EEG PSD relative to baseline (i.e., the most negative value of the β-band SMR modulation curve) from the “Ready” cue to contraction termination (Toriyama *et al*., 2018; Sugino & Ushiyama, 2021), and the magnitude of SRBS (SRBSmax) was defined as the corresponding maximum percentage increase observed during the contraction period. We then quantified the MRBD-to-SRBS transition during voluntary contraction as the difference between SRBS and MRBD (ΔSRBS–MRBD), calculated as SRBSmax − MRBDmax, to capture the shift in neuronal activity from an execution-related desynchronized state to a maintenance-related synchronized state (Fig. 1E, right).

#### CMC analysis

For Experiment 1, 5-s segments extracted from the “Contraction” phase (2–6 s) in all valid trials of each target condition were assembled (24 trials × 5 s = 120 s) to calculate the overall CMC. The EEG and rEMG signals were segmented into 1-s epochs without overlap. A Hanning window was applied to each segment to reduce spectral leakage (Farmer *et al*., 1993; Baker *et al*., 1997; Gross *et al*., 2000). Coherence between EEG and rEMG signals was calculated using the following equation (Halliday *et al*., 1995):

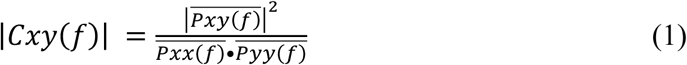

where 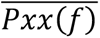 and 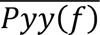 represent the PSD of the EEG and rEMG signals at a given frequency (*f*), respectively, and 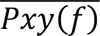 represents the cross-PSD between these two signals. The coherence value *C_xy_*(*f*) ranges from 0 to 1, with 1 indicating complete correlation. To reduce the possibility of coherence values being falsely identified as significant (Type I errors), a Bonferroni correction was applied to adjust the 95% significance level (SL) for multiple comparisons across frequencies (Gwin & Ferris, 2012; Petersen *et al*., 2012; Ushiyama *et al*., 2017):

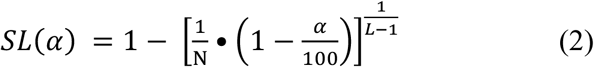

where *L* is the number of non-overlapping 1-s segments used to estimate coherence (i.e., 120 segments per condition in the present study). *N* is the number of frequency bins included in the multiple-comparison correction (i.e., *N* = 48; between 3–50 Hz), and *α* denotes the confidence level (i.e., *α* = 95). Only the maximal values of overall CMC that exceeded the SL were considered significant (CMC+). Participants who exhibited significant CMC in either or both tasks in Experiment 1 were selected to participate in Experiment 2. Because Experiment 2 was designed to compare CMC across task conditions, participants needed to be exhibit detectable CMC in at least one condition, particularly given that β-band CMC is reduced during dynamic force output (Andrykiewicz *et al*., 2007; Omlor *et al*., 2007).

#### Time–Frequency CMC

To calculate CMC in the time series, EEG segmentation was identical to that described for the MRBD analysis. The rEMG signals from each trial were segmented into 1-s epochs with 50-ms sliding steps, following the same procedures as for EEG. For each time point, the 1-s epochs from all 24 trials within the same task condition were concatenated for PSD estimation using Welch’s method (Hanning window; overlap, 0). This procedure produced 360 CMC estimates across the 18-s task period (excluding the 7-s “Rest” phase), based on a data matrix of (1000 data points × 360 windows × 24 trials), which were assembled into a time–frequency CMC map. At each time point, time-resolved β-band CMC was calculated as the integral of the coherence spectrum within 15–35 Hz. The increase in β-band CMC (CMCinc) was defined as the peak value of this β-band CMC curve during the “Contraction” period (Fig. 1F, right).

To quantify the time-resolved co-variation between β-band SMR and CMC, cross-correlation functions (Boker *et al*., 2002) were computed between their modulation curves during the contraction phase (0–7 s). In Experiment 1, data from the 10%- and 15%-tasks were pooled, yielding 55 task observations (10%, n = 28; 15%, n = 27). In Experiment 2, nine CMC+ participants yielding nine observations per task. For each observation in the two experiments, β-band SMR and CMC time courses were z-score normalized, and cross-correlations were calculated across exploratory lags of ±3 s, with positive lags indicating that β-band SMR preceded CMC. Cross-correlation coefficients were Fisher z-transformed (Yang *et al*., 2025) and averaged across observations. Each observation’s peak coefficient was tested against a null distribution generated by shuffling the CMC time series (1,000 permutations). Group-level deviations from zero were assessed across lags using a one-tailed cluster-based permutation test (5,000 sign-flip permutations) (Maris & Oostenveld, 2007). The peak correlation was defined as the maximum value of the group-averaged cross-correlation function.

In addition to the main analysis using the broad β-band, we also performed a narrow-band analysis to account for individual variability in the frequency of maximal CMC as a supplementary analysis, and to enable a more precise comparison between the CMC magnitude and the β-band SMR modulation. We calculated ΔSRBS–MRBD within a 2-Hz frequency band centered on each participant’s peak CMCinc frequency (i.e., ±1 Hz around the peak frequency of the CMCinc peak). This targeted measure was also used for the correlation analysis between CMCinc and ΔSRBS–MRBD. The results showed a similar pattern to those obtained from the broad β-band analysis, as described in the Results section.

The necessity of EMG rectification for CMC analysis remains debated. Indeed, some studies have argued that EMG rectification may be inappropriate (Neto & Christou, 2009; McClelland *et al*., 2012), while others have suggested that rectification is useful (Farina *et al*., 2013; Ward *et al*., 2013). Thus, we additionally conducted CMC analyses using raw EMG (non-rectified EMG) signals and compared the results with those derived from rEMG. The results using raw EMG are provided in the Appendix.

#### Performance Analysis

Force tracking accuracy was quantified by calculating the root-mean-square-error (RMSE) of force output (Waters-Metenier *et al*., 2014) between the recorded force trajectory and the target force trajectory for each trial. To minimize the influence of the initial ballistic contraction deviation and anticipatory relaxation near the end of contraction, force signals were analyzed within 1 to 6 s of the contraction phase. RMSE of force was calculated per trial and averaged across trials to yield a single value per participant per task. Lower RMSE values indicated better force tracking performance.

#### Statistical Analyses

Statistical analyses and data visualization were performed using R software (Version 4.4.3, R Foundation for Statistical Computing, Vienna, Austria) and GraphPad Prism (Version 10, GraphPad Software, Boston, MA, USA). For Experiment 1, because the data were not normally distributed, Wilcoxon signed-rank tests were used to compare CMCinc, MRBDmax, SRBSmax, and ΔSRBS–MRBD between contraction intensities. Associations between CMCinc and ΔSRBS–MRBD, MRBDmax and SRBSmax were assessed using Spearman’s correlation (*ρ*) (α = 0.05), computed separately for each condition. To examine whether behavioral performance was associated with β-band SMR or CMC modulation, Spearman’s correlations were also computed between RMSE of force output and ΔSRBS–MRBD, MRBDmax, SRBSmax and CMCinc. Ninety-five percent confidence intervals (95%CI) for ρ were obtained using a nonparametric bootstrap (10,000 resamples) (Haukoos & Lewis, 2005). Statistical significance was evaluated using a two-sided permutation test (10,000 permutations) (Winkler *et al*., 2014). Scatter plots were overlaid with a Theil-Sen robust trend line for visualization (Wilcox, 1998). Since MRBD is a power decrease and thus negative, ΔSRBS–MRBD can be written as SRBSmax + |MRBDmax|, i.e., the sum of two components for which larger values denote stronger beta modulation. To examine whether and how MRBD and SRBS contributed to the observed association between ΔSRBS–MRBD and CMCinc, partial Spearman correlations (Kim, 2015) were computed, controlling for SRBSmax when assessing the absolute value of MRBDmax (|MRBDmax|), and vice versa. Expressing both components on this common scale makes the signs of the two partial correlations directly comparable. Cross-correlation coefficients at the group-level peak lag were compared between CMC+ and CMC− (non-significant CMC) observations using a linear-mixed model.

Experiment 2 included nine participants who had demonstrated significant CMC between the sensorimotor EEG (Cz) and rEMG from the TA in at least one of the task conditions in Experiment 1. Friedman tests with Dunn’s *post hoc* tests were conducted to evaluate differences in CMCinc, MRBDmax, SRBSmax, ΔSRBS–MRBD, and cross-correlation coefficients at the group-level peak lag across motor tasks (α = 0.05). Association between CMCinc and ΔSRBS–MRBD were assessed using repeated-measures correlation (Bakdash & Marusich, 2017) to quantify the common intra-individual relationship across conditions. In addition, CMCinc values computed from rectified and non-rectified EMG were compared using the Wilcoxon signed-rank test, with the results reported in the Appendix.

## Results

### Experiment 1

#### Inter-Individual Variability in β-Band SMR and CMC Modulation Without a Force-Level Effect

Individual differences were observed in the magnitude of β-band SMR modulation and CMC. Significant CMC between EEG over the sensorimotor area (channel Cz) and rEMG from the TA muscle was observed in nine participants during the 10%-task and in seven participants during the 15%-task; seven participants exhibited significant CMC in both conditions. At the group level, there were no significant differences between the 10%- and 15%-tasks in the magnitudes of CMCinc (10%, 0.089 ± 0.035; 15%, 0.087 ± 0.041; *W* = −81, *p* = 0.315) (Fig. 2A), MRBDmax (10%, −26.65 ± 14.62%; 15%, −28.60 ± 16.04%; *W* = −27, *p* = 0.745) (Fig. 2B), SRBSmax (10%, 24.26 ± 28.41; 15%, 21.19 ± 28.30; *W* = −73, *p* = 0.442) (Fig. 2C) or ΔSRBS–MRBD (10%, 52.64 ± 34.17%; 15%, 52.14 ± 38.15%; *W* = −29, *p* = 0.727) (Fig. 2D) during force output. In addition, CMCinc computed from rEMG was significantly greater than that computed from raw EMG in the 10% task (*W* = −256, *p* = 0.003) (Fig. A1A, left), whereas the difference was not significant in the 15% task (*W* = −102, *p* = 0.229) (Fig. A1B, left).

**Figure 2:**
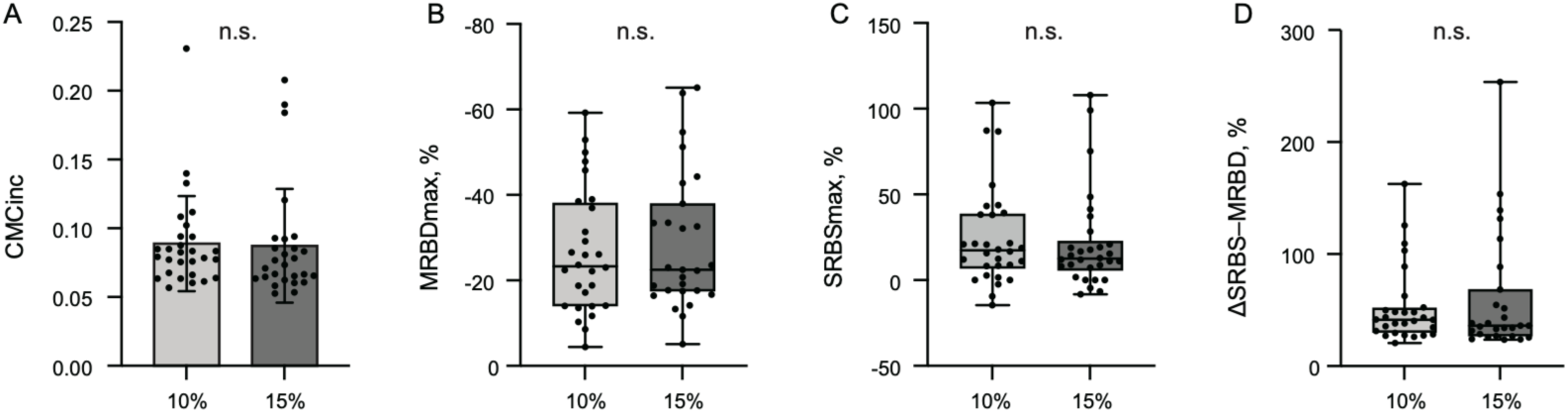
Data for CMC, MRBDmax, SRBSmax and ΔSRBS–MRBD between the 10%-(n = 28) and 15%-tasks (n = 27) in Experiment 1. (A) Group data (mean ± SD) for the CMCinc in the 10%- and 15%-tasks. (B) Boxplots (median, interquartile range, and range) for MRBDmax in the 10%- and 15%-tasks. (C) Boxplots (median, interquartile range, and range) for SRBSmax in the 10%- and 15%-tasks. (D) Boxplots for ΔSRBS–MRBD in the 10%- and 15%-tasks. Light gray bars/boxes represent data from the 10%-task, and dark gray bars/boxes represent data from the 15%-task. Black dots show individual participant data points. “n.s.” indicates no significant differences between the tasks.

#### Concomitant Time-resolved Changes in β-Band SMR Modulation and CMC

Figure 3 illustrates representative examples from four participants, and shows the time-resolved evolution of β-band SMR responses and CMC along with force. Importantly, we observed that the time-resolved evolution of β-band SMR responses over the sensorimotor cortex varied among individuals during motor tasks, and these individual variations were accompanied by corresponding variations in CMC. Participant A displayed a strong MRBD before motor execution, and this was followed by the β-power returning to baseline level during sustained contraction (Fig. 3A). Participant B also showed a prominent MRBD but had a rebound (SRBS) that exceeded baseline levels (Fig. 3B). Participant C demonstrated little MRBD but still exhibited a clear SRBS above the baseline (Fig. 3C). In these participants with relatively greater ΔSRBS–MRBD, CMC emerged at the task onset and diminished toward the end in parallel with the β-power changes (Fig. 3A–C). By contrast, Participant D showed sustained MRBD with minimal ΔSRBS–MRBD and no significant CMC (Fig. 3D).

**Figure 3:**
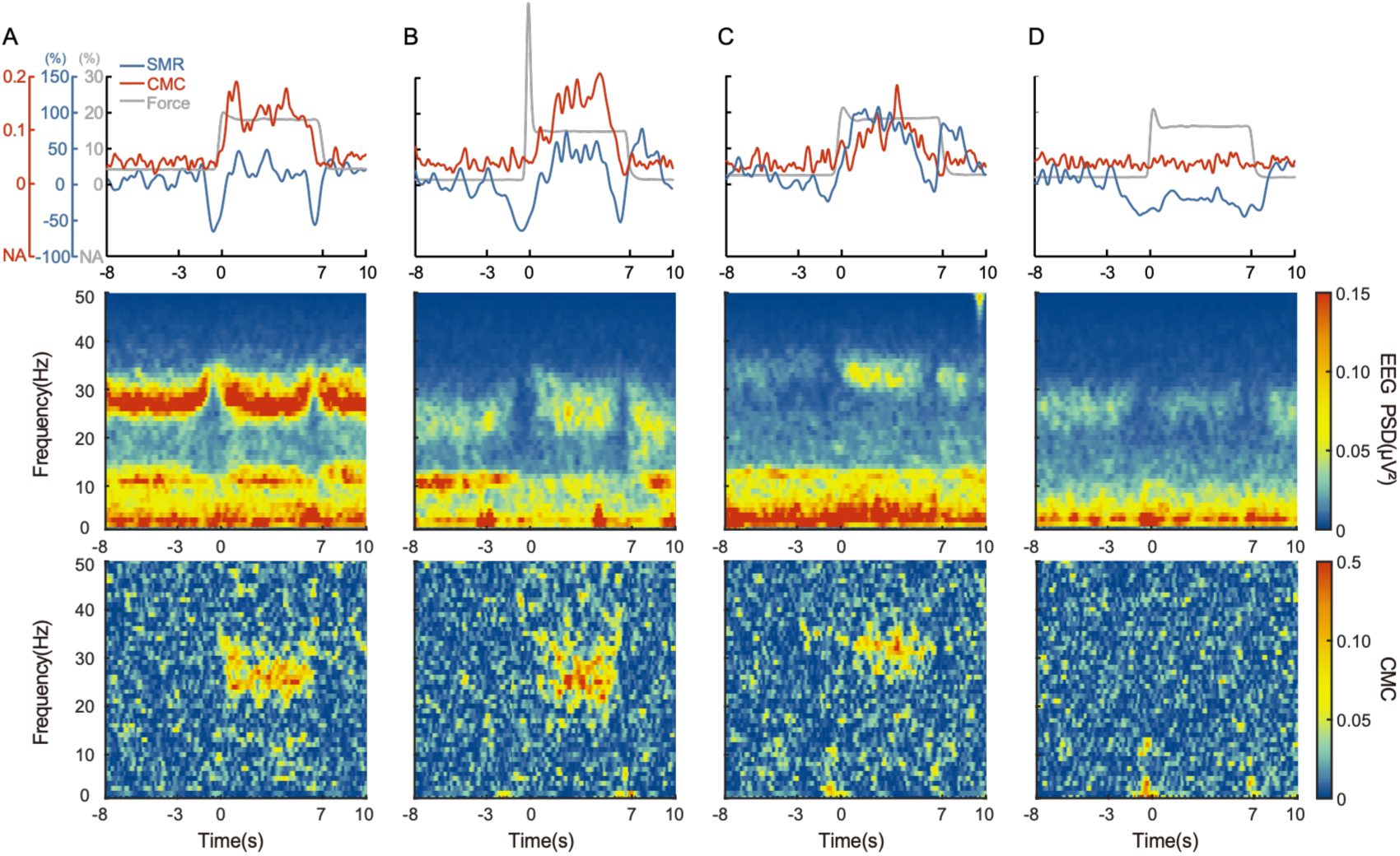
Examples of time-resolved β-band SMR and CMC along with force, EEG time– frequency map, and CMC time–frequency map obtained from four participants in Experiment 1. (A–C) Data from participants with higher CMC values. (D) Data from a participant with no significant CMC. The top row shows dynamic changes in the β-band SMR (blue), CMC (red), and force (gray). The second row exhibits time–frequency maps of EEG power spectral density (PSD) obtained from the Cz channel. The third row shows CMC time–frequency maps.

On the basis of the aforementioned qualitative association between ΔSRBS–MRBD and CMC, we proceeded to quantify this correlation. A positive correlation was identified between ΔSRBS–MRBD and CMCinc in both tasks (10%, *ρ* = 0.611, 95% CI [0.280, 0.830], *p* < 0.001; 15%, *ρ* = 0.690, 95% CI [0.406, 0.859], *p* < 0.001) (Fig. 4A, B). In addition, positive correlations were also observed in SRBSmax and CMCinc in both tasks (10%, *ρ* = 0.500, 95% CI [0.110, 0.781], *p* = 0.008; 15%, *ρ* = 0.510, 95% CI [0.087, 0.801], *p* = 0.007) (Fig. 4C, D), while more negative MRBDmax was associated with greater CMCinc, being non-significant in the 10%-task (*ρ* = −0.290, 95% CI [−0.622, 0.113], *p* = 0.138) (Fig. 4E) but became significant in the 15%-task (*ρ* = −0.540, 95% CI [−0.802, −0.144], *p* = 0.005) (Fig. 4F). To disentangle the respective contributions of MRBD and SRBS, partial Spearman correlations indicated that SRBSmax was positively associated with CMCinc after controlling for |MRBDmax| (10%, *ρ* = 0.546, *p* = 0.003; 15%, *ρ* = 0.529, *p* = 0.005). |MRBDmax| was also positively associated with CMC after controlling for SRBSmax, reaching significance in the 15%-task but not in the 10%-task (10%, *ρ* = 0.381, *p* = 0.0501; 15%, *ρ* = 0.557, *p* = 0.003). These results indicate that CMC reflects contributions from both MRBD and SRBS processes, and is not significantly influenced by either process alone.

Beyond these magnitude-based associations, we further examined the time-resolved dynamics of this co-variation between β-band SMR modulation and CMC. At the individual level, 50 of 55 observations showed positive peak cross-correlations, with mean peak *r* = 0.41 for the 10%-task and *r* = 0.43 for the 15%-task. At the group level, the averaged cross-correlation function showed a significant positive cluster spanning lags from −0.25 to 0.95 (*p* = 0.025; Fig. 4G). The maximum cross-correlation occurred within the cluster at a lag of 0.15 s (*r* = 0.156), indicating that β-band SMR dynamics tended to precede CMC dynamics by approximately 0.15 s. At this group-level peak lag, CMC+ observations (N = 16) showed stronger cross-correlations than CMC− observations (N = 39, *t*_(53.0)_ = 4.365, *p* < 0.001; Fig. 4H). These results demonstrate a reliable time-resolved co-variation between β-band SMR and CMC during sustained contraction, with β-band SMR modulation preceding CMC and is more pronounced when significant CMC is present.

**Figure 4:**
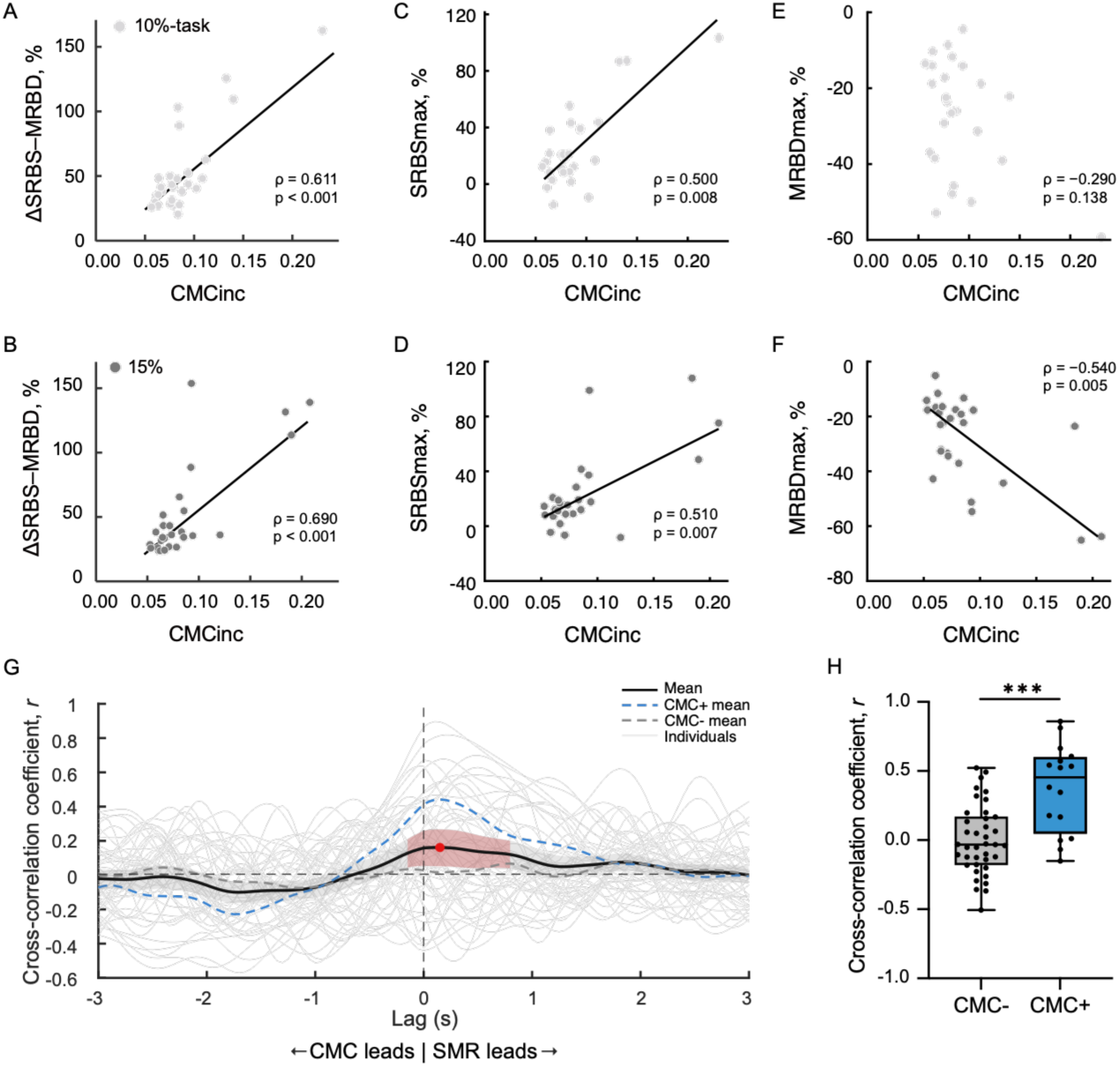
Correlations between the magnitude of CMCinc and ΔSRBS–MRBD, SRBSmax, and MRBDmax of the EEG signal in the 10% (upper row, n = 28), 15%-task (middle row, n = 27), and cross-correlation between time-resolved β-band SMR and CMC (bottom row, n = 55) using data from Experiment 1. (A,B) ΔSRBS–MRBD vs. CMCinc. (C,D) SRBSmax vs. CMCinc. (E,F) MRBDmax vs. CMCinc. Spearman’s correlation coefficient (*ρ*) and permutation test *p*-value are indicated. The black lines represent the Theil-Sen robust trend lines for visualization. The light gray dots represent individual participant data for the 10%-task, and the dark gray dots represent data for the 15%-task. (G) Cross-correlation functions for all observations (N = 55, thin gray lines). The thick black line represents the overall group mean; dashed blue and gray lines indicate the mean for CMC+ and CMC− participants, respectively. Red shaded regions indicate significant lag intervals, and the red dot marks the group peak at 0.15 s. Positive lags indicate that β-band SMR preceded CMC; negative lags indicate that CMC preceded β-band SMR. (H) Comparison of cross-correlation coefficients between β-band SMR and CMC at the peak lag for participants with significant (CMC+) and non-significant (CMC−) CMC, \*\*\**p* < 0.001.

As results of supplementary analyses, the correlation analysis using the 2-Hz frequency band CMC and ΔSRBS–MRBD remained significant (10%, *ρ* = 0.635, 95% CI [0.296, 0.844], *p* < 0.001; 15%, *ρ* = 0.598, 95% CI [0.220, 0.846], *p* = 0.002), which supports the consistency of the observed link between ΔSRBS–MRBD and CMC regardless of inter-individual variability in frequency distribution. In addition, the correlation analysis using raw EMG did not reach statistical significance (10%, *ρ* = 0.122, 95% CI [−0.290, 0.502], *p* = 0.540; Fig. A1A, right; 15%, *ρ* = 0.340, 95% CI [−0.033, 0.635], *p* = 0.074) (Fig. A1B, right).

#### Force-tracking accuracy was not associated with β-band SMR modulation or CMC

Force-tracking accuracy (RMSE) was 1.168 ± 0.772% of MVC for the 10%-task and 1.138 ± 0.802% of MVC for the 15%-task. RMSE was not significantly correlated with MRBDmax (10%, *ρ* = 0.012, 95% CI [-0.333, 0.337], *p* = 0.950; 15%, *ρ* = -0.190, 95% CI [-0.532, 0.195], *p* = 0.343), SRBSmax (10%, *ρ* = -0.105, 95% CI [-0.510, 0.298], *p* = 0.585; 15%, *ρ* = 0.118, 95% CI [-0.300, 0.504], *p* = 0.561), ΔSRBS–MRBD (10%, *ρ* = -0.042, 95% CI [-0.455, 0.341], *p* = 0.831; 15%, *ρ* = 0.199, 95% CI [-0.231, 0.571], *p* = 0.323), or CMCinc (10%, *ρ* = 0.138, 95% CI [-0.261, 0.508], *p* = 0.484; 15%, *ρ* = 0.078, 95% CI [-0.331, 0.499], *p* = 0.696) in either condition, indicating that inter-individual differences in β-band SMR modulation and CMC were not explained by differences in force-tracking performance.

### Experiment 2

Individual variability was observed in Experiment 1. β-band SMR activity in some participants appeared to return to a near-resting state, indicating a rebound of β-band SMR power (SRBS). Conversely, some participants exhibited a sustained reduction in β-band SMR during steady force maintenance. These distinct patterns, together with the observed significant correlations, raise the possibility that the presence and strength of CMC may be determined by how β-band SMR activity is modulated in the process of steady force maintenance. On the basis of these findings, we hypothesized that if tasks require more complex force adjustment, the simple rhythmic control that supports steady contraction may not fully meet the task demands. In such cases, rhythmic β-band SMR activity might be disrupted in accordance with task requirements, so that even individuals who exhibited greater CMC during steady force maintenance in Experiment 1 might shift toward reduced β-band power rebound (i.e., continuous MRBD), accompanied by weaker CMC. To explore whether β-band SMR modulation and CMC change as a function of task type (i.e., steady-force maintenance vs. continuous force modulation) or in parallel with increasing task difficulty, we conducted Experiment 2 using varying task lines that induced different levels of force adjustment difficulty.

#### Concomitant Changes in β-Band SMR Modulation and CMC Across Tasks

Figure 5 illustrates group-averaged data across all participants for each task in Experiment 2. Similar to the findings in Experiment 1, the modulation of CMC coincided with the β-power rebound following MRBD during muscle contraction in all four tasks of Experiment 2. However, the magnitude of these changes varied depending on the task type. The task that required steady force maintenance (the St-task) exhibited stronger SRBS and CMC, whereas the tasks involving continuous force adjustments (the 1sin-, 3sin-, and 4sin-tasks) tended to show weaker SRBS and reduced CMC.

**Figure 5:**
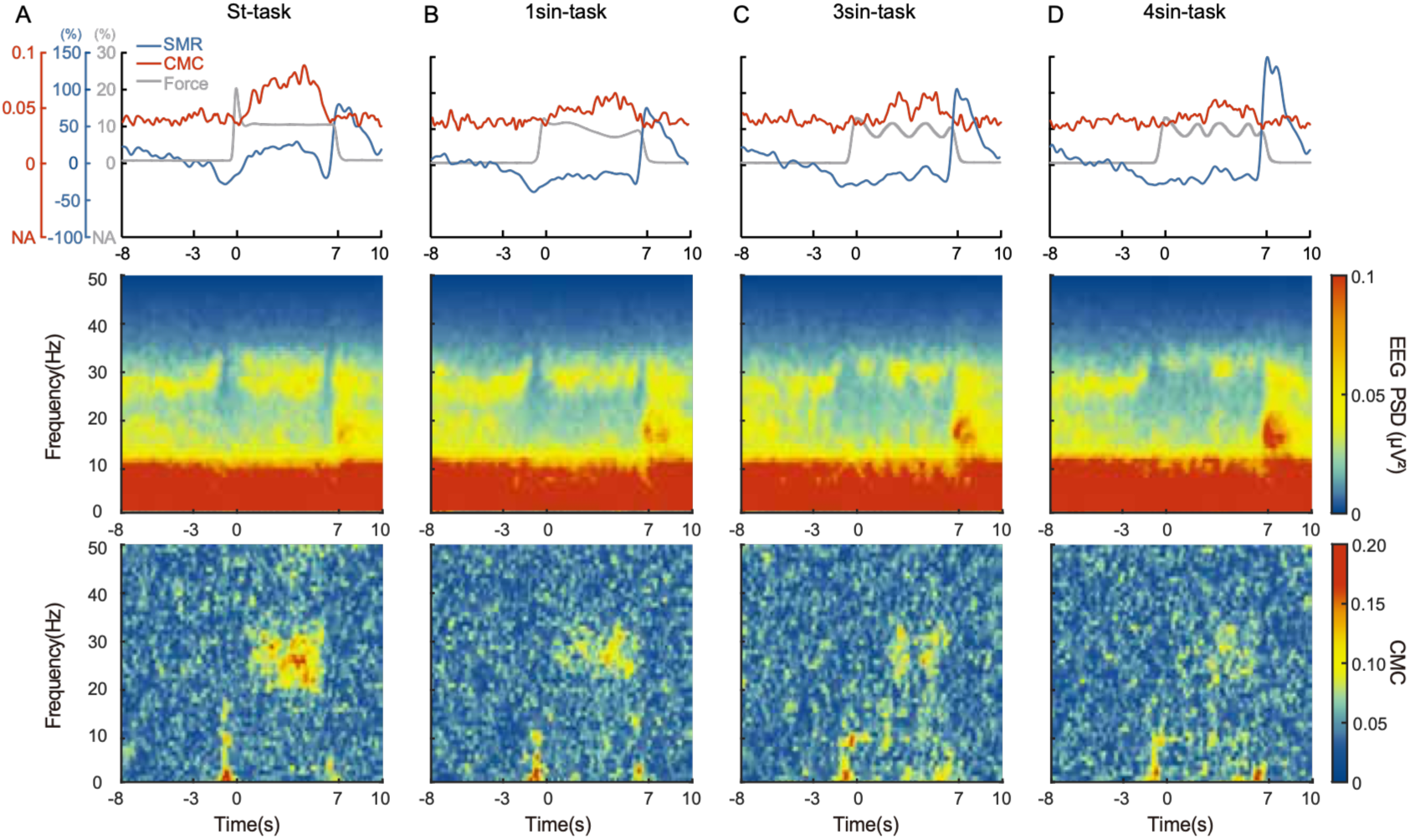
Group-averages of the time-resolved β-band SMR and CMC along with torque, EEG time–frequency maps, and CMC time–frequency maps for each task in Experiment 2 (n = 9). (A) Data from St-task. (B) Data from 1sin-task. (C) Data from 3sin-task. (D) Data from 4sin-task. The top row shows dynamic changes in β-band SMR (blue), CMC (red), and force (gray). The second row shows the time–frequency maps of EEG PSD for the Cz channel. The third row presents the CMC time–frequency maps.

#### Task-Dependent Intra-Individual Variability in β-Band SMR and CMC Modulation

Figure 6A and B show the grand average of time-resolved CMC and β-band SMR response from all participants for each task, respectively. Both time-resolved CMC and β-band SMR response during the contraction phase exhibited greater increases in the St-task than in the other three tasks. By contrast, the three curved-line tasks (the 1sin, 3sin, and 4sin-tasks) exhibited similar patterns, with no substantial differences among them. Figure 6C–F show the group data for the CMCinc, ΔSRBS–MRBD, SRBSmax, and MRBDmax during muscle contraction, respectively, under the four task conditions.

**Figure 6:**
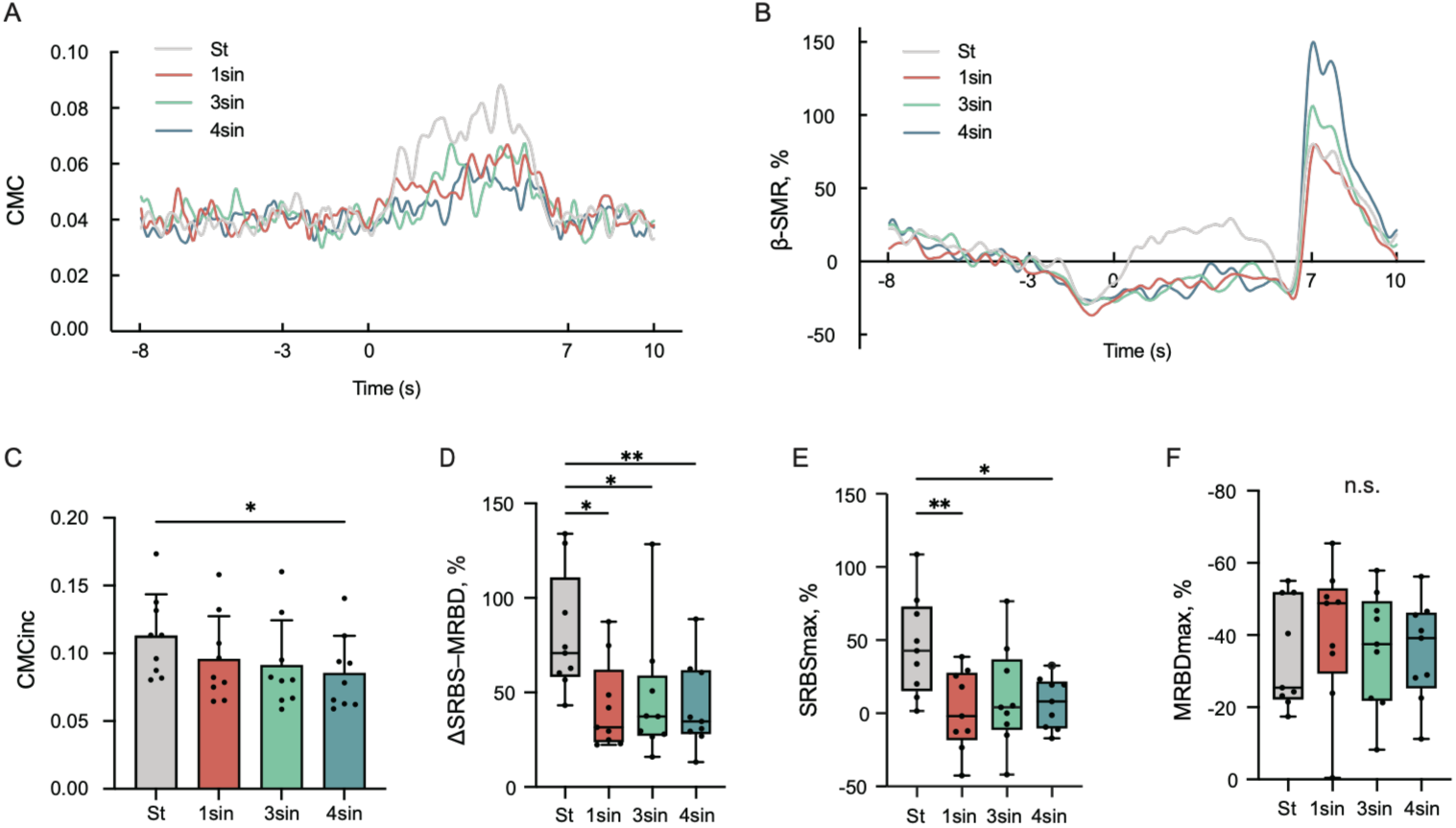
Group data for Experiment 2 (n = 9). (A) Grand average of time-resolved β-band CMC across all participants for the St-task (gray), 1sin-task (red), 3sin-task (green), and 4sin-task (blue). (B) Grand average of time-resolved β-band SMR across all participants for the St-task (gray), 1sin-task (red), 3sin-task (green), and 4sin-task (blue). (C) Group data (mean ± SD) for CMCinc across the four tasks. (D,E,F) Boxplots (median, interquartile range, and range) showing the distribution of individual data for ΔSRBS– MRBD (D), SRBSmax (E) and MRBDmax (F) across the four tasks; black dots represent individual participant data. \**p* < 0.05, \*\**p* < 0.01.

The Friedman test revealed a significant effect of task condition on CMCinc (χ^2^(3) = 10.20, *W* = 0.38, *p* = 0.017), ΔSRBS–MRBD (χ^2^(3) = 14.20, *W* = 0.53, *p* = 0.003) and SRBSmax (χ^2^(3) = 13.40, *W* = 0.50, *p* = 0.004). Multiple comparisons revealed that the CMCinc in the St-task was significantly greater than that in the 4sin-task (*Z* = 3.10, *p* = 0.012). The ΔSRBS–MRBD was also significantly greater in the St-task than in the other three tasks, whereas no significant differences were observed among the 1sin-, 3sin-, and 4sin-tasks (St-task vs. 1sin-task, *Z* = 3.10, *p* = 0.012; St-task vs. 3sin-task, *Z* = 2.74, *p* = 0.037; St-task vs. 4sin-task, *Z* = 3.29, *p* = 0.006). In addition, the SRBSmax in the St-task was significantly greater than that in the 1sin and 4sin-tasks (St-task vs. 1sin-task, *Z* = 3.47, *p* = 0.003; St-task vs. 4sin-task, *Z* = 2.74, *p* = 0.037). Conversely, there was no significant difference in the magnitude of MRBDmax across the tasks (χ^2^ (3) = 0.20, *W* = 0.01, *p* = 0.978).

For supplementary analyses, when both CMCinc and ΔSRBS–MRBD were computed within the 2-Hz frequency bands, the results replicated the main findings: there were significant task effects for ΔSRBS–MRBD (χ^2^ (3) = 8.60, *W* = 0.44, *p* = 0.008) and CMCinc (χ^2^ (3) = 11.80, *W* = 0.32, *p* = 0.035). *Post ho*c tests revealed that ΔSRBS– MRBD in the St-task was significantly greater than that in all other tasks (St-task vs. 1sin-task: *Z* = 2.92, *p* = 0.021; St-task vs. 3sin-task: *Z* = 2.74, *p* = 0.037; St-task vs. 4sin-task: *Z* = 2.74, *p* = 0.037), whereas CMCinc in the St-task was significantly greater than that in the 4sin-task (*Z* = 2.74, *p* = 0.037), thereby mirroring the results from the broad β-band analysis. In addition, CMCinc did not differ significantly between rEMG and raw EMG estimates in any task conditions (St-task, *W* = 5.00, *p* = 0.820; 1sin-task, *W* = −15.00, *p* = 0.426; 3sin-task, *W* = 11.00, *p* = 0.570; 4sin-task, *W* = 23.00, *p* = 0.203) (Fig. A2A). CMCinc computed using raw EMG also differed across tasks, showing a significant main effect of task (χ^2^ (3) = 9.80, *W* = 0.36, *p* = 0.020), although none of the pairwise comparisons reached significance, suggesting a trend that was consistent with the main findings.

We next analyzed the repeated measure correlation between ΔSRBS–MRBD and CMCinc and identified a strong intra-individual positive relationship (*r*_rm_ = 0.749, df = 26, 95% CI [0.522, 0.877], *p* < 0.001) (Fig. 7A). A similar positive correlation was also observed using the 2-Hz frequency band (*r*_rm_ = 0.483, df = 26, 95% CI [0.134, 0.725], *p* = 0.009), thus supporting the robustness of this relationship. Additionally, when CMCinc derived from raw EMG signals was used, the repeated measure correlation with ΔSRBS–MRBD remained a positive trend but did not reach significance (*r*_rm_ = 0.226, df = 26, 95% CI [−0.042, 0.630], *p* = 0.080) (Fig. A2B).

Finally, we examined the time-resolved dynamics of the co-variation between β-band SMR modulation and CMC across the four tasks. A cluster-based permutation test (one-tailed, r > 0) revealed a significant positive cluster spanning lags from −0.30 to 0.95 s only in the St-task (p = 0.022), while none of the three sinusoidal tasks yielded a significant cluster (Fig. 7B). In St-task, the group-averaged cross-correlation peaked at a lag of 0.35 s (*r* = 0.431), indicating that β-band SMR fluctuations preceded CMC by approximately 0.35 s. The peak lags of the other tasks also occurred at positive lags but their peak correlations were substantially weaker (1sin: 0.30 s, *r* = 0.147; 3sin: 0.20 s, *r* = 0.177; 4sin: 0.20 s, *r* = 0.193). Cross-correlation strength at the task-specific peak lags differed significantly across tasks (χ^2^(3) = 10.73, *W* = 0.40, *p* = 0.013; Fig. 7C); *post hoc* comparisons showed stronger correlations in the St-task than in the 1sin-task and 3sin-task (St-task vs. 1sin-task: *Z* = 2.92, p = 0.021; St-task vs. 3sin-task: *Z* = 2.74, p = 0.037). Together, these results indicate that the time-resolved co-variation between β-band SMR and CMC was strongest during steady maintenance contraction, with SMR preceding CMC, and was attenuated when the task required continuous force adjustment.

**Figure 7:**
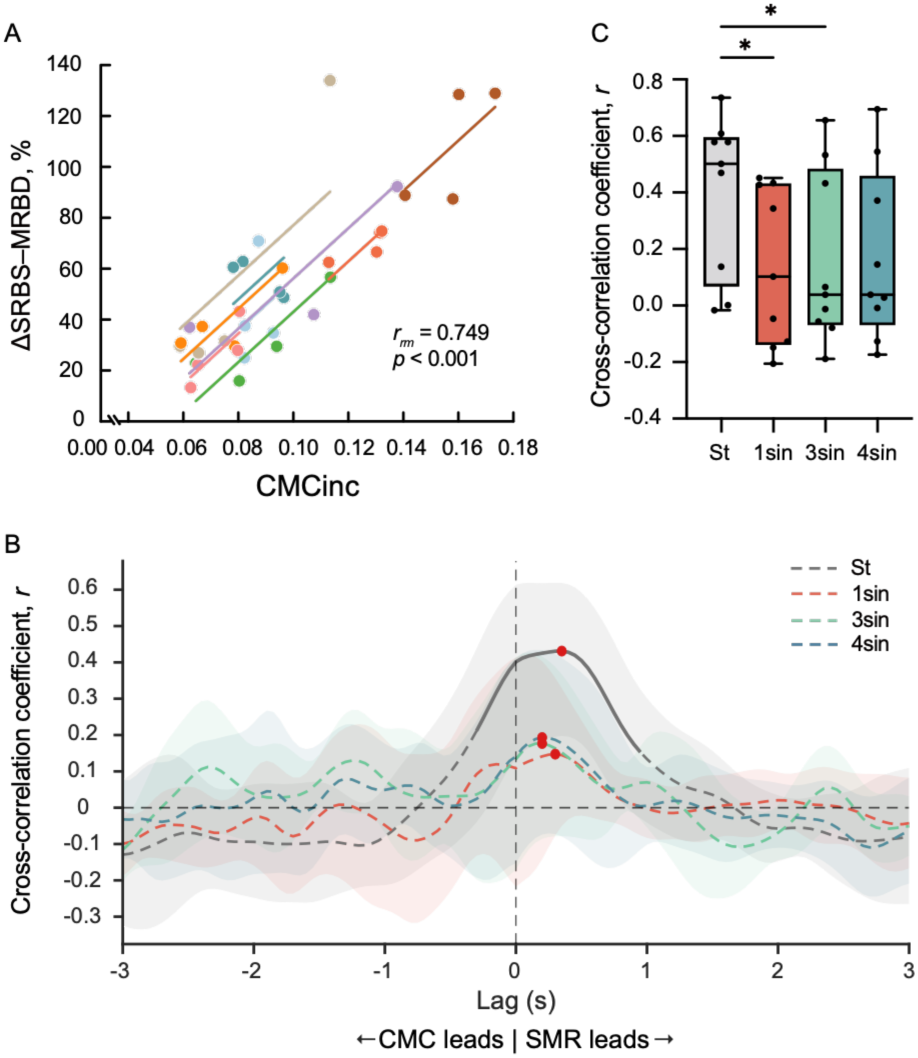
Relationship between β-band CMC and SMR modulation in Experiment 2 (n = 9). (A) Repeated measure correlation between the magnitude of CMCinc and ΔSRBS–MRBD of the EEG signal across the four task conditions in Experiment 2. The same participant is given the same color, with corresponding lines to show the repeated measure correlation fit for each participant. Repeated measure correlation coefficient (*r_rm_*) and the *p*-value are indicated. (B) Cross-correlation functions between β-band SMR and CMC for the St-task (gray), 1sin-task (red), 3sin-task (green), and 4sin-task (blue); shaded regions indicate the 95% confidence interval. Solid line segments indicate significant lag intervals, and red dots mark each task’s peak lag. Positive lags indicate that β-band SMR preceded CMC, while negative lags indicate that CMC preceded β-band SMR. (C) Boxplots (median, interquartile range, and range) showing the distribution of individual cross-correlation coefficients at the task-specific peak lags across the four tasks; black dots represent individual participant data. \**p* < 0.05.

#### Force tracking performance confirms differential task difficulty

To verify whether the task manipulation produced different levels of force-tracking difficulty in Experiment 2, RMSE was calculated between the recorded and target force trajectories for each task condition. The Friedman test revealed a significant effect of task condition on RMSE (χ^2^(3) = 21.13, *W* = 0.78, *p* < 0.001), demonstrating that force-tracking difficulty differed significantly across the four tasks. Multiple comparisons showed that the 4sin-task produced significantly higher RMSE than the St-task (*Z* = 4.02, *p* < 0.001), and 1sin-task (*Z* = 3.83, *p* < 0.001). No significant differences were found among the St-, 1sin-, and 3sin-tasks (St vs. 1sin: *p* = 1.000; St vs. 3sin: *p* = 0.268; 1sin vs. 3sin: *p* = 0.407), nor between the 3sin- and 4sin-tasks (3sin vs. 4sin: *p* = 0.268). These results indicate that the 4sin-task was associated with the greatest tracking error, being significantly higher than the St- and 1sin-tasks, thereby confirming that the task manipulation induced meaningful differences in behavioral difficulty.

## Discussion

To address how β-band SMR modulation relates to CMC, we examined two experimental contexts: inter-individually during steady-force maintenance and intra-individually during dynamic-force adjustment (Fig. 1). Our main findings were as follows. First, in participants with stronger CMC, the time-resolved enhancement of CMC coincided with the EEG β-power rebound from MRBD to SRBS during steady-force contraction (Fig. 3). The ΔSRBS–MRBD varied across individuals and was positively correlated with CMC magnitude (Fig. 4A-F), with both MRBD and SRBS contributing to this association. Cross-correlation analysis further characterized this time-resolved co-variation, revealing that β-band SMR modulation preceded CMC, with this leading relationship being significantly more pronounced in participants with significant CMC (Fig. 4G, H). These neural associations were independent of force-tracking accuracy across individuals. Second, intra-individual comparisons across tasks revealed that both ΔSRBS–MRBD and CMC were concurrently reduced in tasks required continuous force adjustment (Fig. 6). However, increasing task difficulty within the continuous-adjustment condition had no comparable effect. Across all task conditions, the two measures remained positively correlated intra-individuals (Fig. 7A). Cross-correlation analysis further confirmed that this time-resolvedly structured co-variation (with SMR preceding CMC) was strongest during steady-force maintenance and weakened under continuous force adjustment (Fig. 7B, C). Together, these results indicate that the extent of β-band SMR modulation is associated with the strength of CMC (i.e., the two co-vary across individuals and are modulated within individuals depending on task demands).

### Experiment 1: Inter-Individual Differences in β-Band SMR Modulation and CMC

In Experiment 1, β-band SMR modulation during steady-force maintenance aligned with changes in CMC. Specifically, individuals who showed a greater ΔSRBS–MRBD tended to exhibit stronger CMC, whereas those who remained in a persistent MRBD state without subsequent β-power rebound exhibited weaker CMC. These results corroborate and integrate previous findings of a positive correlation between CMC and EEG β-power (Kristeva *et al*., 2007; Ushiyama *et al*., 2011, 2017), as well as the positive association between CMC and the magnitude of MRBD (Bayraktaroglu *et al*., 2013; Sun *et al*., 2025). While these findings may appear contradictory, our results reconcile them by quantifying the full modulation range of the β-band SMR during steady contraction. Specifically, individual differences in CMC are associated with the full β-band SMR modulation profile from MRBD to SRBS, rather than being limited to the absolute EEG β-power during contraction. This is critical because individuals with comparable absolute EEG β-power during contraction may exhibit fundamentally different β-band SMR modulation trajectories. Indeed, ΔSRBS–MRBD captures information that static power metrics cannot access, reflecting contributions from both β-power suppression and rebound processes.

We showed that the absolute strength of MRBD positively correlated with CMC, indicating that stronger attenuation of β-oscillation around contraction onset related to greater CMC. Previous studies have proposed that stronger initial MRBD reflects increased cortical excitability and a breakdown of local spatial synchronization, which in turn might facilitate the subsequent recruitment of neuronal networks responsible for establishing reliable corticospinal control, as reflected in greater CMC (Bayraktaroglu *et al*., 2013). Our findings complement this interpretation. Although stronger initial MRBD was associated with greater CMC, we observed that when MRBD persisted throughout the steady contraction without a subsequent resynchronization, CMC tended to be weak. In contrast, the restoration of local spatial synchronization after MRBD may reflect the active recruitment of cortical networks involved in maintaining the current motor set (Engel & Fries, 2010). In addition, a recent motor-unit-based neurofeedback study showed that participants could volitionally up- or down-modulate β-activity in both peripheral and cortical signals, but β-band CMC did not differ between up and down conditions (Bräcklein *et al*., 2022). This implies that CMC is not simply determined by absolute β-power. Accordingly, we propose that the β-band SMR transition from MRBD to SRBS, as quantified by ΔSRBS–MRBD, serves as a functional correlate of CMC, rather than explained solely by MRBD or by β-power itself.

Beyond these magnitude-based associations, cross-correlation analysis further quantified the time-resolved co-variation between β-band SMR and CMC, indicating that β-band SMR modulation tended to precede CMC by approximately 0.15 s during steady contraction in the Experiment 1. Given the limited time-resolved resolution of this analysis and the inherent imprecision of scalp EEG in resolving detailed cortical processes, this finding should be interpreted as a trend rather than a precise time-resolved estimate. Nevertheless, the cluster suggests that the emergence of corticospinal coupling may follow the EEG β-band power rebound after MRBD, a pattern that was particularly evident in participants with significant CMC. This time-resolved precedence is biologically plausible, given that peripheral β-band oscillations are driven by cortical β-band activity transmitted via the fastest corticospinal fibers (Abbagnano *et al*., 2025), implying that cortical β-band changes would necessarily precede the emergence of CMC. To our knowledge, this is the first study to reveal the time-resolved precedence of β-band SMR modulation over CMC during steady contraction using a time-resolved analysis approach.

Although individual differences in MRBD and PMBR have been widely studied (Parkes *et al*., 2006b; Espenhahn *et al*., 2017b; Saha & Baumert, 2020), SRBS following MRBD has received less attention. Previous studies have primarily focused on group-averaged tendencies (Cassim *et al*., 2000; Alegre *et al*., 2006; Riddle & Baker, 2006) and have rarely consider how this pattern might vary across individuals. Our findings indicate that not all participants exhibit increased EEG β-power during steady-force maintenance. Such inter-individual variability reflects different propensities for synchronized or desynchronized modes of neuronal activity during steady-force maintenance.

In the Experiment 1, RMSE of force output was not significantly correlated with CMC or β-band SMR modulation in either condition. This finding is consistent with previous research reporting no significant correlation between CMC and motor output steadiness or accuracy within or across individuals (Perez *et al*., 2006; Johnson *et al*., 2011; Larsen *et al*., 2017), but appears to contrast with studies reporting a significant association between CMC and force control accuracy (Kristeva *et al*., 2007; Mendez-Balbuena *et al*., 2012; Ushiyama *et al*., 2017). One possible explanation for this discrepancy is that the relatively low task difficulty resulted in uniformly good performance across participants, limiting the behavioral variance necessary to detect a correlation with neural measures. More broadly, this pattern is consistent with the view that β-band oscillations are not essential for task performance per se, but rather reflect a form of neural control that the motor system employs to varying extents across individuals (Kilner *et al*., 1999; Perez *et al*., 2006).

### Experiment 2: Intra-Individual Differences in β-Band SMR Modulation and CMC

Experiment 2 examined whether and how the association between β-band SMR modulation and CMC would be changed in relation to force adjustment difficulty using intra-individual approach. Although we obtained significant difference in RMSE of force output among tasks, the changes in ΔSRBS–MRBD and CMC did not scale with these performance differences. In contrast, both ΔSRBS–MRBD and CMC were significantly reduced during dynamic-force adjustment tasks compared with steady-force conditions (St-task). These results suggest that these neural measures are sensitive to the contraction type (i.e., whether the task required steady-force maintenance or continuous force adjustment) rather than to the degree of task difficulty. Moreover, a positive intra-individual correlation indicated that task-dependent changes in β-band SMR co-varied with corticospinal coupling strength, further suggesting that ΔSRBS– MRBD serves as a functional correlate of CMC across different task demands. Previous findings support this task-dependent changes. Indeed, EEG β-power quickly returns to baseline after the motor initiation during sustained wrist extension (Cassim *et al*., 2000), whereas MRBD persists without recovery during repetitive finger movements (Erbil & Ungan, 2007). These contrasting patterns suggest that the descending motor command required to maintain a steady force output differs from that required to adjust the force output. While simple force maintenance may benefit from synchronized neural activity, this neural communication mode dominated by a single β-rhythm may be too simplistic for the demands of continuous force adjustment. Thus, complex force adjustment appears to require desynchronized neural activity, as reflected in reduced ΔSRBS– MRBD and CMC.

The dynamic-force adjustment tasks differed from the simple steady-force maintenance task by eliciting continuous MRBD rather than SRBS during contraction. Although manipulation of the task parameters successfully altered task difficulty, as reflected by differences in force-tracking accuracy quantified by RMSE, neither MRBD nor CMC differed significantly across the three dynamic-force adjustment tasks. This is consistent with previous findings that task parameters such as kinematics and kinetics have a negligible effect on MRBD (Stancák & Pfurtscheller, 1996; Pistohl *et al*., 2012; Nakayashiki *et al*., 2014), and that CMC does not differ across several low-frequency force modulation conditions within the 0–2 Hz range (Naranjo *et al*., 2010). Our research extended these findings by showing that MRBD magnitude remained consistent across different speeds of the dynamic force adjustment task, suggesting a relatively constant neural drive available to generate CMC irrespective of task difficulty. Cross-correlation analysis further revealed that the time-resolved relationship between β-band SMR modulation and CMC was also task-dependent. The lead–lag relationship observed in Experiment 1 was reproduced in the St-task of Experiment 2, whereas peak cross-correlation coefficients were markedly reduced during the dynamic force-adjustment tasks. The reduced cross-correlation suggests that continuous force adjustment weakens the stable temporal coordination between cortical β-band activity and CMC. Such a task may require ongoing updating of motor commands, which could prevent the sensorimotor system from maintaining the relatively stable rhythmic β-band activity observed during steady-force maintenance.

Although not a main focus of our investigation, all tasks showed pronounced PMBR. Previous studies have shown that stronger PMBR is associated with greater movement speed and acceleration, suggesting that it may reflect an increased need for inhibitory control to stabilize the sensorimotor system following more demanding motor output (Fry *et al*., 2016; Zhang *et al*., 2020). Consistent with this concept, PMBR increased with the number of sine cycles (St < 1sin < 3sin < 4sin), indicating that increased demands for frequent and rapid force adjustments evoke stronger post-movement inhibitory responses. The PMBR may serve as a functional index of neural inhibitory processes to stabilize the sensorimotor system after motor output.

### Integration of Experiments 1 and 2: Functional Significance of β-Band SMR Co-Variation

The inter-individual differences in Experiment 1, ranging from persistent MRBD with insignificant CMC to pronounced SRBS with strong CMC, may reflect a diversity of control strategies across individuals. Some participants exhibited EEG β-band power increase during steady-state contraction, which has been interpreted as a reliance on cortical synchronization to maintain the current motor state and resist changes (Engel & Fries, 2010; Matta *et al*., 2025). This was accompanied by stronger CMC, which has been proposed to support more efficient corticospinal communication by optimally timing mutual inputs between the cortex and spinal motoneurons (Schoffelen *et al*., 2005; Mendez-Balbuena *et al*., 2012). By contrast, some participants appeared not to exhibit EEG β-band power increase during steady-force maintenance, as indicated by sustained MRBD. This is considered to reflect increased cortical activation in task-relevant sensorimotor areas (Pfurtscheller & Lopes da Silva, 1999; Neuper & Pfurtscheller, 2001). Persistent desynchronization may present a more effortful, top-down control mode in which the cortex continuously monitors and adjusts motor output even under steady task demands, with reduced β-band oscillatory coupling caused by desynchronized motor neuron activity. Our findings suggest that individuals may engage distinct degrees of cortical and corticomuscular synchronization even under identical task conditions, and might flexibly modulate the extent of both depending on task demands.

Previous studies support our interpretation that neural strategies can be flexibly adjusted, with β-band SMR modulation and its downstream expression in CMC changing in parallel. CMC can be strengthened through task-specific motor training (Pohja & Salenius, 2003; Mendez-Balbuena *et al*., 2012), suggesting a shift toward a more efficient strategy characterized by a stronger β-power rebound (SRBS) and CMC. Conversely, motor imagery elicits β-band ERD (Pfurtscheller *et al*., 2005; Duann & Chiou, 2016)—particularly when imagining finger tapping during static force holding (Nijhuis *et al*., 2021)—which reflects a shift toward a more effortful strategy characterized by sustained β-power suppression (MRBD). Together, these findings suggest that concurrent β-band SMR and CMC modulation reflects flexible adjustments in synchronization-based neural strategies, which are shaped by task demands and individual neural propensity.

### Limitations

A potential technical limitation of our study is that surface EEG may fail to capture β-band SMR activity in some participants, possibly because of biological factors such as corticospinal neuron orientation or skull and cerebrospinal fluid properties (Malmivuo *et al*., 1997; Olejniczak, 2006). Indeed, several participants in Experiment 1 showed neither evident MRBD nor significant CMC during muscle contraction, possibly because of detection limitations. However, among other participants without significant CMC, EEG β-band activity was successfully recorded. Specifically, a subset of participants exhibited clear MRBD without subsequent SRBS, whereas the remaining participants showed a small degree of MRBD and ΔSRBS–MRBD. These patterns indicate that an absence of significant CMC was not primarily caused by a failure to detect β-band activity but rather reflected insufficient neural resynchronization during contraction. Thus, although technical issues cannot be completely excluded, we believe that they did not substantially affect the observed relationship between CMC magnitude and the extent of the β-band SMR transition. Moreover, only participants with significant CMC were included in Experiment 2, further minimizing the potential influence of such issues. A further consideration concerns the relatively lower proportion of significant CMC individuals (9 of 30) in our participants than that reported in previous studies (e.g., Ushiyama *et al*., 2011). Such a difference would be expected given that CMC has been shown to be weaker during contractions lasting only a few seconds than during prolonged sustained ones (Suzuki & Ushiyama, 2020). This suggests that CMC may be engaged to varying extents depending on task demands, reflecting a flexibility in neural control within individuals.

Some studies have argued that EMG rectification is not necessary for CMC analysis because this causes distortion of EMG signal (McClelland *et al*., 2012) whereas other works indicate that oscillatory input components may be strengthened by rectification, particularly at low contraction levels (Farina et al., 2013).Because of this debate, we also report CMC computed from raw EMG (non-rectified EMG) in the Appendix to ensure transparent reporting. As shown in Fig. A1, CMC estimated from raw EMG was not consistent with that derived from rEMG, particularly in the 10%-task, in which CMCinc was significantly reduced. Though the degree of discrepancy varied across individuals, the reductions in CMC values due to non-rectification of EMG were particularly pronounced in participants with greater CMC values estimated using rEMG. These findings are consistent with the view that coherence computed from raw EMG may fail to reveal a detectable peak at the input frequency, particularly at low contraction levels (Farina *et al*., 2013). In the present study, the small-amplitude EMG may not accurately capture grouped discharges without rectification at low contraction levels, thereby resulting in an underestimation of CMC and no significant correlation between ΔSRBS–MRBD and CMCinc. Although no general conclusion can be drawn about the necessity of rectification, rectification process of EMG may provide a more accurate estimate of CMC, at least within the present protocol. The source of the inter-individual variability in the impact of rectification needs further investigation.

## Conclusion

Overall, we clarified the relationship between β-band SMR modulation and CMC by examining inter-individual differences during steady-force maintenance and intra-individual variations across tasks with varying demands. In this context, ΔSRBS– MRBD, which incorporates contributions from both neural desynchronization and resynchronization processes, may serve as a functional correlate of CMC at both the individual level and across task demands. Even under identical steady-force conditions, individuals adopted different patterns of cortical regulation and CMC, thus reflecting individual propensities to engage in a synchronization-based neural strategy. Additionally, steady-force maintenance was associated with a synchronization-based strategy with greater SRBS and CMC, whereas dynamic-force adjustment favored a desynchronization-based strategy with persistent MRBD and weaker CMC. Collectively, these findings indicate that the way in which the nervous system regulates β-band SMR may serve as a strategy for flexible adaptation to diverse motor environments.

## Appendix

Figure A1, A2

**Figure A1.**
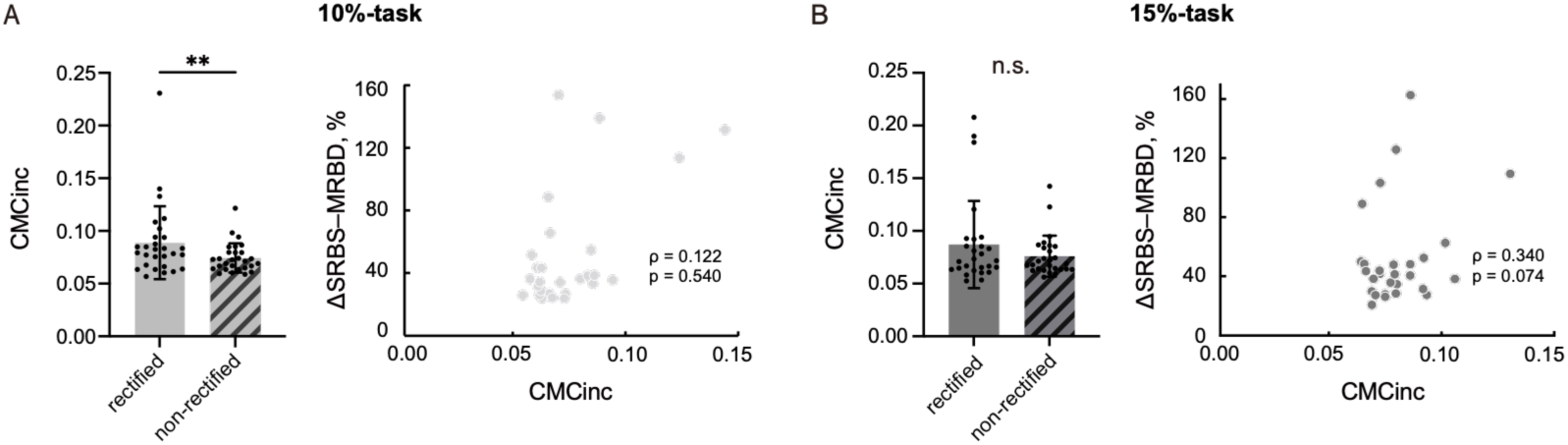
Results from Experiment 1 using raw EMG. (A) 10%-task (n = 28) and (B) 15%-task (n = 27). Left: Comparison between CMCinc computed from rectified (solid) and non-rectified (hatched) EMG (mean ± SD). Black dots indicate individual participant data points. “n.s.” indicates not significant, and ** indicates *p* < 0.01. Right: Spearman correlations between ΔSRBS–MRBD and CMCinc computed using raw EMG; *ρ* and permutation-test *p*-values are shown.

**Figure A2:**
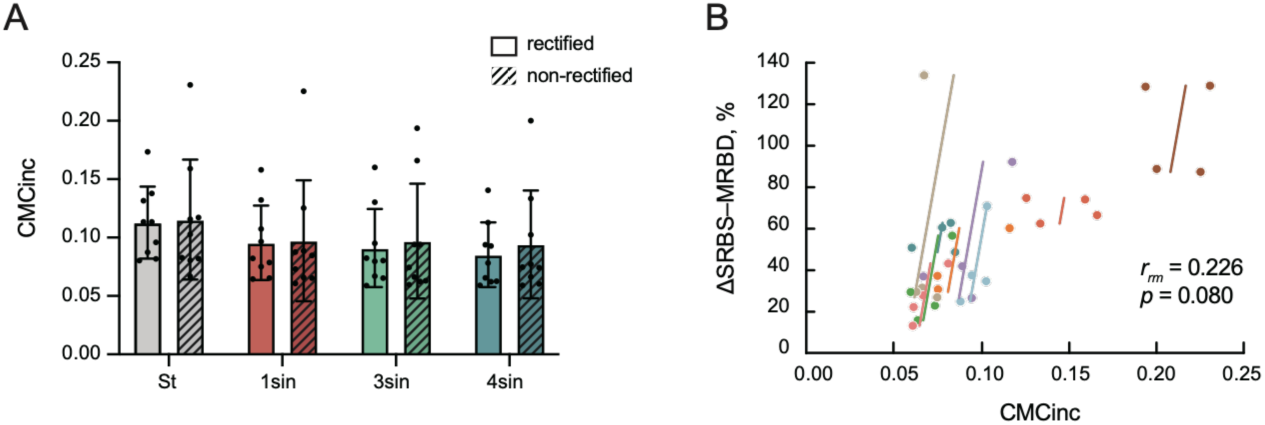
Results using raw EMG in Experiment 2 (n = 9). (A) Group data (mean ± SD) for CMCinc in the St-, 1sin-, 3sin-, and 4sin-tasks, computed from rectified (solid) and non-rectified (hatched) EMG. Black dots indicate individual participant data points. (B) Repeated measure correlation between the magnitude of CMCinc computed using raw EMG and ΔSRBS–MRBD of the EEG signal across the four task conditions. The same participant is given the same color, with corresponding lines to show the repeated measure correlation fit for each participant. The repeated measure correlation coefficient (*r_rm_*) and the *p*-value are indicated.

## Acknowledgements

We thank Ms. Tomomi Hamaoka for her secretarial assistance, Mr. Takuya Ideriha and Mr. Hirotaka Sugino for their help with data analyses, and all other members of our laboratory for their helpful discussions. We thank Bronwen Gardner, PhD, from Edanz (https://jp.edanz.com/ac) for editing a draft of this manuscript. A preprint version of this manuscript has been deposited on bioRxiv (https://doi.org/10.1101/2025.11.10.687531).

